# csppinet: A Python package for context-specific biological network construction and analysis based on omics data

**DOI:** 10.1101/2023.05.23.541999

**Authors:** L.M. de Carvalho

## Abstract

The analysis of large-scale biological data, particularly omics data, has become essential for understanding complex biological systems. In this study, we present a Python package for the construction of context-specific biological networks based on gene expression activity and protein-protein interaction (PPI) data. The package leverages computational tools and the NetworkX library for network analysis. Through a case study focusing on yeast fermentation with glucose and xylose as carbon sources, we demonstrate the package’s capabilities. The context-specific networks derived from these fermentation conditions were compared to highlight the impact of different carbon sources and stages on network dynamics. Hub genes were identified within each network, and Gene Ontology (GO) and Kyoto Encyclopedia of Genes and Genomes (KEGG) analyses were performed to understand their functional implications. The results revealed distinct hub genes and enriched biological processes in each context-specific network. In the glucose-specific network, the exclusively proteins in this network revealed enrichment in terms related to *chromosome organization, biological regulation, MAPK signaling pathway* and *mismatch repair pathway*. Conversely, the exclusively proteins in xylose-specific network revealed enrichment in *generation of precursor metabolites and energy), mitochondrion organization, response to extracellular stimulus, glutamate metabolic process, cellular response to alcohol, beta-Alanine metabolism, arginine and proline metabolism, glyoxylate and dicarboxylate metabolism* and *pyruvate metabolism*. The developed Python package and context-specific networks provide researchers with a valuable framework to explore complex biological phenomena. By integrating gene expression profiles and PPI data, researchers can gain insights into context-dependent molecular interactions and regulatory mechanisms. These findings contribute to the understanding of cellular behavior and have potential implications in disease mechanisms, biomarker identification, and drug target discovery. The csppinet package is available at https://github.com/lmigueel/csppinet.

## 1. Introduction

In the era of genomics and personalized medicine, the analysis of large-scale biological data has become increasingly vital for understanding complex biological systems. Among the various types of high-throughput data, omics data holds a prominent position as it provides a comprehensive snapshot of cellular processes at different molecular levels. However, deciphering the intricate relationships between genes, proteins, and other molecular entities hidden within these vast datasets presents a significant challenge.

One of the fundamental concepts in systems biology is the protein-protein interaction (PPI) network, which offers a holistic view of how proteins interact and collaborate within a cell. PPI networks provide a valuable framework for unraveling the intricate web of molecular interactions underlying cellular functions and are essential for studying disease mechanisms, drug discovery, and understanding biological processes at a systems level. By exploring the topology, connectivity, and functional associations of proteins, researchers can gain insights into complex biological phenomena and identify key molecular players driving specific cellular processes.

While traditional PPI networks serve as valuable references, they are often generic and fail to capture the dynamic nature of biological systems. Cells are highly adaptable entities, and their behavior can vary depending on the environmental conditions, developmental stages, or disease states they experience. To address this limitation, the concept of context-specific networks has emerged, aiming to generate customized networks that reflect the specific cellular context under investigation [1-3].

Context-specific networks are constructed by integrating omics data, such as gene expression activity, with the existing knowledge of PPI networks [4]. By incorporating context-specific information, researchers can gain a deeper understanding of the functional interactions and regulatory mechanisms that govern cellular processes within a specific context. These networks enable the identification of context-dependent biomarkers, uncover novel drug targets, and provide insights into the molecular mechanisms underlying complex diseases.

In this article, we introduce a novel Python package designed to facilitate the construction and analysis of context-specific biological networks based on omics data. By integrating gene expression profiles, PPI databases, and mathematical algorithms, our package offers an easy-to-use object-oriented Python API and a command-line interface (CLI) for context-specific network construction and post-analysis.

## 2. Implementation

### 2.1 Construction

To implement our proposed Python package for generating context-specific biological networks, we utilized a modular and extensible approach. The package was developed using Python programming language, leveraging various libraries and tools to facilitate efficient network construction and analysis. Our pipeline is shown in Figure 1. The construction of the package involved several key components:

**Figure 1.**
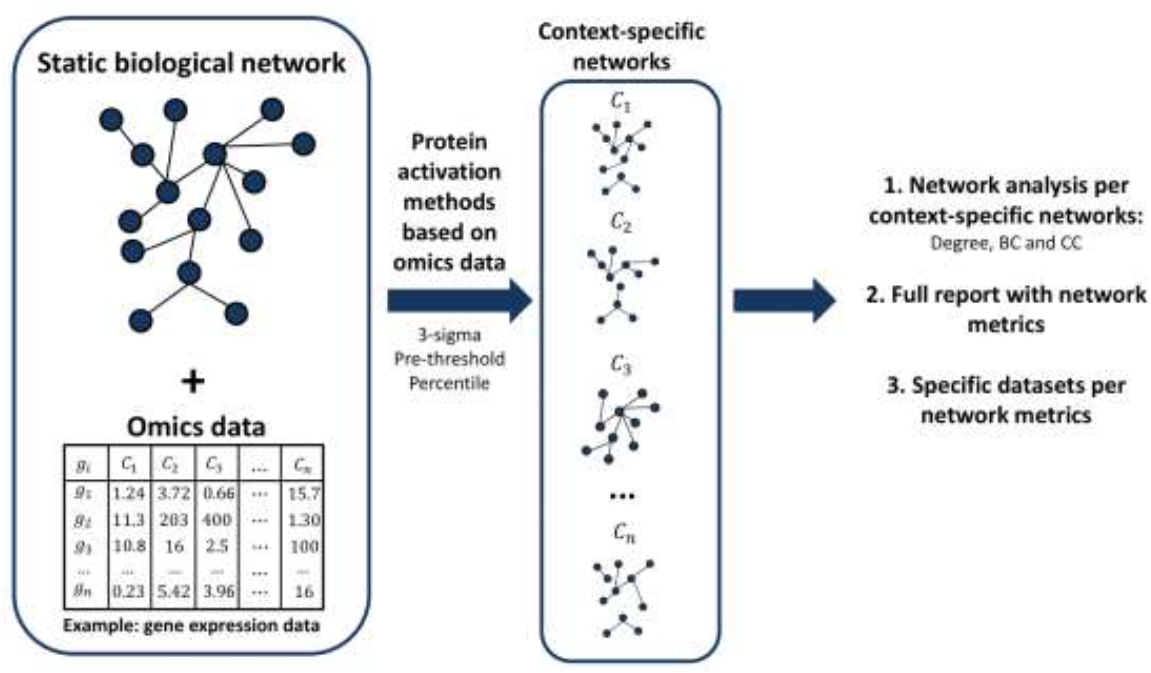
Pipeline for generating context-specific biological networks by csppinet package. The figure illustrates the step-by-step process of generating context-specific biological networks using our proposed pipeline. The pipeline incorporates omics data and protein-protein interaction (PPI) networks or other static biological networks to construct networks that capture the dynamic nature of biological systems. It highlights the key stages, including data preprocessing, integration of gene expression activity, identification of active proteins, and network construction. The pipeline provides a comprehensive framework for contextual analysis, enabling the exploration of specific biological conditions and their impact on network dynamics.

a. Data Integration: The package provides functionality to seamlessly integrate omics data, including gene expression profiles and protein-protein interaction (PPI) networks. The input data, such as expression data in standard formats (e.g., CSV, TSV) and PPI network files (e.g., CSV, edge-list), can be easily loaded and processed.
b. Context-Specific Network Generation: The core functionality of the package lies in the generation of context-specific networks based on gene expression activity. By integrating gene expression profiles with the PPI network, the package identifies expressed proteins at different biological conditions and constructs sub-networks capturing the dynamic interactions between these proteins.
c. Network Analysis: The package incorporates the powerful NetworkX library for network analysis and provides a range of network metrics and algorithms. This allows users to investigate various network properties, such as centrality measures, clustering coefficients, and module identification, providing deeper insights into the structural and functional characteristics of the generated context-specific networks.

### 2.2 Identification of expressed proteins

The implementation of the context-specific biological network package relies on the three-sigma method described by Wang, J et al., 2013 [5], which involves determining the activity of proteins based on gene expression values and constructing context-specific subnetworks; the method proposed by the Tang, X et al., 2011 [6], in which gene expression values are evaluated against a predefined threshold; and the percentile method, where the percentiles of gene expression values is established as a threshold for determining protein activation in each biological condition. This section expands the idea of these methods.

First and foremost, it is important to emphasize that the “3-sigma” method is modeled based on the statistical properties of a normal distribution, which aligns well with datasets that exhibit a larger sample size. As the number of biological samples increases, the “3-sigma” method becomes more effective in accurately identifying active proteins and constructing context-specific biological networks. If the user has a limited number of samples, it is advisable to consider employing the other two methods rather than the “3-sigma” method. This recommendation is based on the observation that when the dataset contains only a few samples, the networks generated by the “3-sigma” method tend to be smaller in size. In such cases, the statistical robustness and efficiency of the “3-sigma” method may not be fully utilized due to the limited amount of data available. Instead, the other methods can offer viable alternatives, as they may be better suited for generating context-specific biological networks with the given dataset. By selecting an appropriate method based on the number of samples, researchers can ensure the optimal utilization of their data and obtain more meaningful insights into protein activation dynamics.

#### 2.2.1 Three-sigma method

Protein activity is crucial for cellular functions, and its regulation involves controlling protein abundance and lifetime within the cell. To deduce protein activity, the method focuses on determining the active time points based on gene expression data. The three-sigma method is employed to differentiate between inactive and active time points during the cellular cycle. This method considers the characteristic expression curves of individual genes and accounts for the inherent noise in gene expression arrays. By designing gene-specific thresholds using the three-sigma principle, it is essential to accurately identify the time points or biological conditions with the highest expression values as the active periods for each protein. This approach provides a comprehensive understanding of protein activity by taking into account individual gene dynamics and the variability in gene expression data.

Also, it is important to highlight that the three-sigma method offers a practical solution to determine protein activity based on gene expression data. It addresses the challenges in accurately identifying active time points by considering the variations in gene expression curves and the presence of noise in the data. By establishing gene-specific thresholds using the three-sigma principle, it is effectively distinguishing between inactive and active time points during the cellular cycle. This approach enables a more precise assessment of protein activity, contributing to a deeper understanding of cellular dynamics and the regulatory mechanisms that control protein abundance and lifetime.

The first step in the implementation involves processing the omics data, such as gene expression profiles, and calculating the gene expression thresholds (Th(p)) for individual proteins. This step utilizes Python’s data processing capabilities and numerical libraries, such as NumPy and Pandas, to efficiently handle large-scale data sets. In order to identify active proteins within a dynamic protein-protein interaction (PPI) subnetwork, considering that *E*_*j*_(*p*) is the gene expression of gene p in the biological condition j, the active threshold Th(p) is:

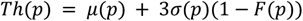

where 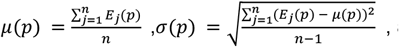, and 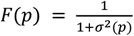.

An active gene/protein is defined as one whose gene expression value is greater than Th(p).

In contrast, a static PPI network is typically represented as an undirected graph, denoted as G(V,E). Here, V denotes the set of nodes representing proteins, while E represents the set of edges representing connections between pairs of proteins, *p*_*i*_ and. *p*_*j*_ To capture the dynamic nature of biological conditions, a PPI subnetwork, *G*_*j*_ (*V*_*i*_,*E*_*j*_) is introduced. Each subnetwork *G*_*j*_ corresponds to a specific biological condition, denoted as *b*_*j*_, with i ranging from 1 to n. In the context-specific network, *p*_*i*_ protein is considered to interact with protein *p*_*j*_ only if they are active in the same biological condition (*b*_*j*_) and if there is a connection between them in the static network. This criterion ensures that protein-protein interactions are restricted to those occurring within a specific biological context, emphasizing the importance of both activity and connectivity in the network.

#### 2.2.2 Based on a predefined threshold

The algorithm based on a predefined threshold to determine a protein expression integrates gene expression profiles with static protein-protein interaction (PPI) networks to generate context-specific protein interaction networks. The algorithm involves two main stages:

1. Identification of expressed proteins: At each biological condition, the expression values of genes are compared against a predefined threshold. If the expression level of a protein exceeds the threshold, it is considered as expressed at that specific biological condition.
2. Construction of sub-networks: Interactions between proteins are considered expressed when both interacting proteins are expressed simultaneously at a particular biological condition. Expressed proteins and their corresponding interactions are extracted from the static network, iteratively creating a series of sub-networks at different biological conditions. These sub-networks collectively represent the dynamic nature of protein interactions in different context-specific conditions.

#### 2.2.3 Percentiles

To determine protein activation levels based on gene expression, we employed the concept of percentiles. Gene expression data obtained from high-throughput technologies often contain noise and variability. By calculating the percentiles of gene expression values, we can establish a threshold for determining protein activation. We chose to focus on a specific percentile value, such as the 5^th^ percentile or the 25th percentile, which represents the expression level above which a protein is considered active. This approach allows us to account for variations in gene expression across different biological conditions and provides a standardized criterion for defining protein activation.

### 2.3 Network analysis

By incorporating these considerations in above sections, the construction of context-specific networks enables a more nuanced analysis of protein interactions, taking into account the dynamic nature of gene expression and the underlying static network topology.

To handle analysis of networks, the package utilizes the popular network analysis library, NetworkX v2.5.1 [7]. NetworkX provides a comprehensive set of tools and algorithms for the manipulation, visualization, and analysis of complex networks. By leveraging the capabilities of NetworkX, the package enables users to perform various network report metrics and analyses on the generated context-specific biological networks.

We used three commonly used metrics for network analysis are degree, closeness centrality, and betweenness centrality. With the intersection of top genes in these lists, the user could identify hub genes.

The node’s degree in a network corresponds to the count of its connections with other nodes. It provides a measure of the node’s importance and involvement in the network. Nodes with higher degrees often indicate crucial hubs or highly connected proteins within the network. Closeness centrality measures how easily information can flow from a given node to all other nodes in the network. Nodes with higher closeness centrality are characterized by their ability to quickly disseminate information to distant parts of the network. This metric is particularly useful in identifying proteins that have a central role in the network and are involved in efficient communication and coordination processes. Betweenness centrality quantifies the extent to which a node serves as a bridge or intermediary for information flow between other nodes in the network. Nodes with high betweenness centrality act as critical connectors, controlling the transfer of information between different regions of the network. These nodes often occupy key positions in pathways or regulatory cascades, playing essential roles in information transfer and facilitating communication between distinct modules or functional units.

## 3. Case of study

To demonstrate the effectiveness of the ccsppinet package, a case study was conducted using data from the STRING database, encompassing all yeast interactions. The study focused on analyzing the gene expression data obtained during a 2G ethanol fermentation process, specifically considering four time points [8]. By comparing the context-specific networks derived from glucose (time point - 7h) and xylose (time point - 16h) consumption, the study aimed to highlight the impact of different fermentation carbon sources and stages on network dynamics. We used the “percentile” method with a threshold of 25 (meaning of 25th percentile or first quartile). Also, we filtered the interactions in the STRING database with “combined_score” > 900. All materials and scripts used in this analysis are available in the Supplementary Material.

Within the context-specific network generated from glucose fermentation, numerous hub genes have emerged as pivotal players in the cellular response to this carbon source. Most notably, a significant portion of these hub genes are ribosomal proteins, contributing to their high-density subnetwork. The glucose-specific network has 3,604 nodes and 51,211 edges, generating 57 components. Given the high number of components, it is possible to identify gene clusters that may possess distinct roles at this stage of fermentation. In contrast, the context-specific network derived from xylose fermentation exhibited distinct hub genes and functional implications. The xylose-specific network has 3,545 nodes and 49,445 edges, generating 56 components. Like glucose-specific network, a significant portion of these hub genes are ribosomal proteins.

To gain further insights into the differences in functional implications of the glucose-specific and xylose-specific networks, we conducted a comprehensive Gene Ontology (GO) and Kyoto Encyclopedia of Genes and Genomes (KEGG) analysis. This analysis focused on proteins that were exclusively present in one network and absent in the other. Remarkably, we discovered 354 unique proteins in the glucose-specific network and 295 unique proteins in the xylose-specific network. Additionally, our investigation revealed a substantial overlap, with 3,250 active proteins found to be shared between the two networks.

The GO analysis over the 354 unique proteins in the glucose-specific network revealed enrichment in terms related to Chromosome organization (GO:0051276; FDR < 0.05), Signaling (GO:0023052; FDR < 0.05) and Biological regulation (GO:0065007; FDR < 0.05). Moreover, the KEGG analysis over these exclusively proteins highlighted the involvement of the genes in pathways such as MAPK signaling pathway (sce04011; FDR < 0.05) and mismatch repair (sce03430; FDR < 0.01), further emphasizing their crucial roles in glucose fermentation.

To explore the functional significance of the 295 unique proteins in the xylose-specific network, GO and KEGG analyses were performed on genes. The GO analysis revealed enrichment in terms related to Generation of precursor metabolites and energy (GO:0006091; FDR < 0.05), Mitochondrion organization (GO:0007005; FDR <0.05), Response to extracellular stimulus (GO:0009991; FDR < 0.05), Glutamate metabolic process (GO:0006536; FDR < 0.05) and Cellular response to alcohol (GO:0097306; FDR < 0.05), reflecting the network’s emphasis on xylose utilization and the cellular response to oxidative stress. Moreover, the KEGG analysis over these unique proteins highlighted the involvement of hub genes in pathways such as beta-Alanine metabolism (sce00410; FDR < 0.05), Arginine and proline metabolism (sce00330; FDR < 0.05), Glyoxylate and dicarboxylate metabolism (sce00630; FDR < 005) and Pyruvate metabolism (sce00620; FDR < 0.05).

Overall, the analysis of the context-specific networks derived from glucose and xylose fermentation provided valuable insights into the key hub genes and functional implications associated with different carbon sources and stages of fermentation. By focusing on these networks and performing GO and KEGG analyses on the genes, the study uncovered the intricate molecular interactions and biological processes underlying glucose and xylose utilization during 2G ethanol fermentation. These findings demonstrate the potential of the context-specific biological network package in unraveling complex biological phenomena and facilitating targeted investigations in the field of systems biology.

## 4. Conclusions

In this article, we present the csppinet package, a Python package for context-specific biological network construction and analysis based on omics data. Here, we also highlighted that the development and utilization of context-specific biological networks based on omics data hold great promise in advancing our understanding of complex biological systems.

Through the implementation of the package, we showcased its ability to generate context-specific networks by integrating gene expression profiles and the existing knowledge of PPI networks. We emphasized the dynamic nature of cellular behavior and highlighted the importance of considering both activity thresholds and network connectivity in constructing context-specific networks.

Furthermore, we presented a case study focused on yeast fermentation with different carbon sources, specifically glucose and xylose. By analyzing the context-specific networks derived from these fermentation conditions, we identified hub genes and performed GO and KEGG analyzes to gain insights into their functional implications.

In conclusion, the development of context-specific biological networks based on omics data offers a valuable approach to unraveling the complexity of biological systems. By considering the dynamic gene expression activity and integrating it with protein-protein interaction data, we can gain insights into context-dependent molecular interactions and regulatory mechanisms. This can lead to a deeper understanding of disease mechanisms, identification of biomarkers, and discovery of potential drug targets.

