## Supplementary material for "csppinet: A Python package for context-specific biological network construction and analysis based on omics data": script_csppinet_STRING.py

```
from urllib import request
import pandas as pd
import csppinet

# The download link for the yeast interactome in TSV format from STRING
url = 'https://stringdb-static.org/download/protein.links.v11.5/4932.protein.links.v11.5.txt.gz'

# Load the interactome data from the URL into a Pandas DataFrame
ppi_data = pd.read_csv(url, sep='\s+', compression='gzip')
ppi_data = ppi_data.replace({'4932.': ''}, regex=True)
ppi_data = ppi_data[ppi_data['combined_score'] > 900]
ppi_data.to_csv("STRING_Yeast_interactions_900_full.csv",index=False)

# csppinet arguments. Input files should be in .csv extension 
network = 'STRING_Yeast_interactions_900_full.csv'
exp = 'full_expression.csv'
threads = 10
method = "percentile"
#method = "3-sigma"
#method = "pre-threshold"
value_percentile = 25 
#value_pthreshold = 5

#construction
csppinet.construction(network,exp,threads,method,value_percentile)

#network metrics
csppinet.network_metrics(exp)
```
